# Antibody-based signal amplification for single-cell proteomics by mass spectrometry

**DOI:** 10.64898/2025.12.22.695898

**Authors:** Jakob Woessmann, Judit Pina Agullet, Anne Louise Blomberg, Bo T. Porse, Steffen Goletz, Erwin M. Schoof

## Abstract

Single-cell proteomics by mass spectrometry (scp-MS) detects thousands of proteins per cell, yet low-abundant proteins routinely detected by antibody-based methods are challenging to quantify in the single-cell proteome. To address this, we developed antibody peptide tags (AbTags) - MS detectable tandem peptide barcodes coupled to antibodies which provide signal amplification, enable absolute quantification, and are quantified alongside endogenous proteomes by scp-MS. As proof-of-concept, AbTags quantify cytokine-induced PDL1 expression, undetectable by scp-MS.

## Main

Mass spectrometry (MS)-based proteomics is a central technology for analyzing protein dynamics across diverse sample types. Recent advances in single-cell proteomics by MS (scp-MS) allows the detection and quantification of thousands of proteins per cell [1], [2]. However, many proteins that are readily detected by antibody-based methods are still missing in scp-MS datasets. Therefore, an external readout such as flow cytometry is often needed to quantify these markers, but many scp-MS workflows do not routinely include such parallel antibody measurements. Conversely, targeted antibody-based assays such as flow cytometry and CyTOF offer highly sensitive quantification of tens of proteins, but provide no information about the accompanying global proteome [3], [4]. Moreover, these methodologies are constrained in multiplexing and instrumentation requirements, and do not allow for absolute quantification of protein targets (i.e. molecule counting). In contrast, RNA sequencing technologies such as CITE-seq combine transcriptome-wide RNA profiling with targeted protein detection via antibody-conjugated oligonucleotide barcodes, which are detected during sequencing [5]. Conceptually analogous LC-MS workflows that couple deep, unbiased single-cell proteomes with an antibody-defined panel of otherwise undetectable protein targets in the same cell are currently missing.

Here, we introduce antibody peptide tags (AbTags), tandem-repeat peptide barcodes conjugated to antibodies that are read out alongside the endogenous single-cell proteome by LC-MS (**Fig. 1A**). AbTags consist of short, protease-releasable peptide repeats that are attached to established antibody backbones as fusion proteins at either the N- or C-terminus via a linker. During standard scp-MS sample preparation, digestion releases multiple copies of each barcode per antigen-binding event, providing signal amplification in defined numbers and decoupling assay sensitivity from endogenous protein abundance. Each AbTag carries a unique peptide sequence, analogous to distinct fluorochromes in flow cytometry, enabling multiplexed panels whose theoretical scale exceeds that of existing antibody-based cytometry methods. As a proof-of-concept, we implemented the AbTag concept towards two proteins, PDL1 and HER2, which we could not quantify endogenously from single DU-145 prostate cancer cells by data-independent acquisition (DIA) scp-MS. Two well-established antibodies (IgGs against PDL1 and HER2) [6] were selected and conjugated either C-or N-terminally with a range of tandem peptide repeats (1x, 8x and 16x) via a flexible 10-mer glycine-serine (GGGGS)_2_ linker to the heavy chain, resulting in 2x, 16x, and 32x peptide repeats per AbTag molecule, respectively (**Fig. 1** **C-D, Supplementary Fig. 1**). Using wide-window parallel reaction monitoring (wwPRM) and DIA [7], we demonstrate that AbTag peptides are digested and quantified with high precision (CV 2.2 %, n = 8, **Supplementary Fig. 2E**) and accuracy across at least three orders of magnitude (**Fig. 1B**). Furthermore, by comparing AbTags with 2- and 32-peptide repeats, we confirm the expected signal amplification due to the increased number of peptide repeats, leading to a 16-fold increase in dilution points above Limit of Detection (LOD). While we observe minor decreases in binding efficiency of the AbTag, scaling with the number of peptide-repeats, binding capacity similar to control IgG’s could be achieved at relevant labeling concentrations (**Fig. 1C-D**). Next, through actual scp-MS experiments (**Fig. 1E**), we stained DU-145 cells with both the anti-PDL1 and anti-HER2 AbTags. Peptide-repeat based signal amplification (>2x) was required to quantify the AbTag peptide in scp-MS (**Fig. 1** **F-G**). FACS-based AbTag detection did not show a significant difference between the PDL1 and HER2 expression of the cells stained with different AbTags (**Supplementary Fig. 2F, H**). Quantitative linearity between FACS-based AbTag- and LC-MS AbTag detection could be observed for wwPRM based peptide barcode quantification (PDL1 and HER2), as well as for DIA based peptide barcode quantification (only PDL1) (**Fig. 1H, Supplementary Fig. 2G**, I, J). Having established the quantitative robustness of AbTag, we next evaluated the ability to leverage absolute quantitation abilities afforded by stable-isotope-labeled (SIL) peptides matching the AbTag peptide sequences. Thereby we were able to report the absolute number of antibody binding events per cell (median 42,683 anti-PDL1 molecules, 7,299 anti-HER2 molecules) (**Fig. 1I**) which represents a powerful proxy for absolute antigen expression on target cells and demonstrates the ability to conduct absolute quantitation with antibody-based technologies.

**Figure 1.**
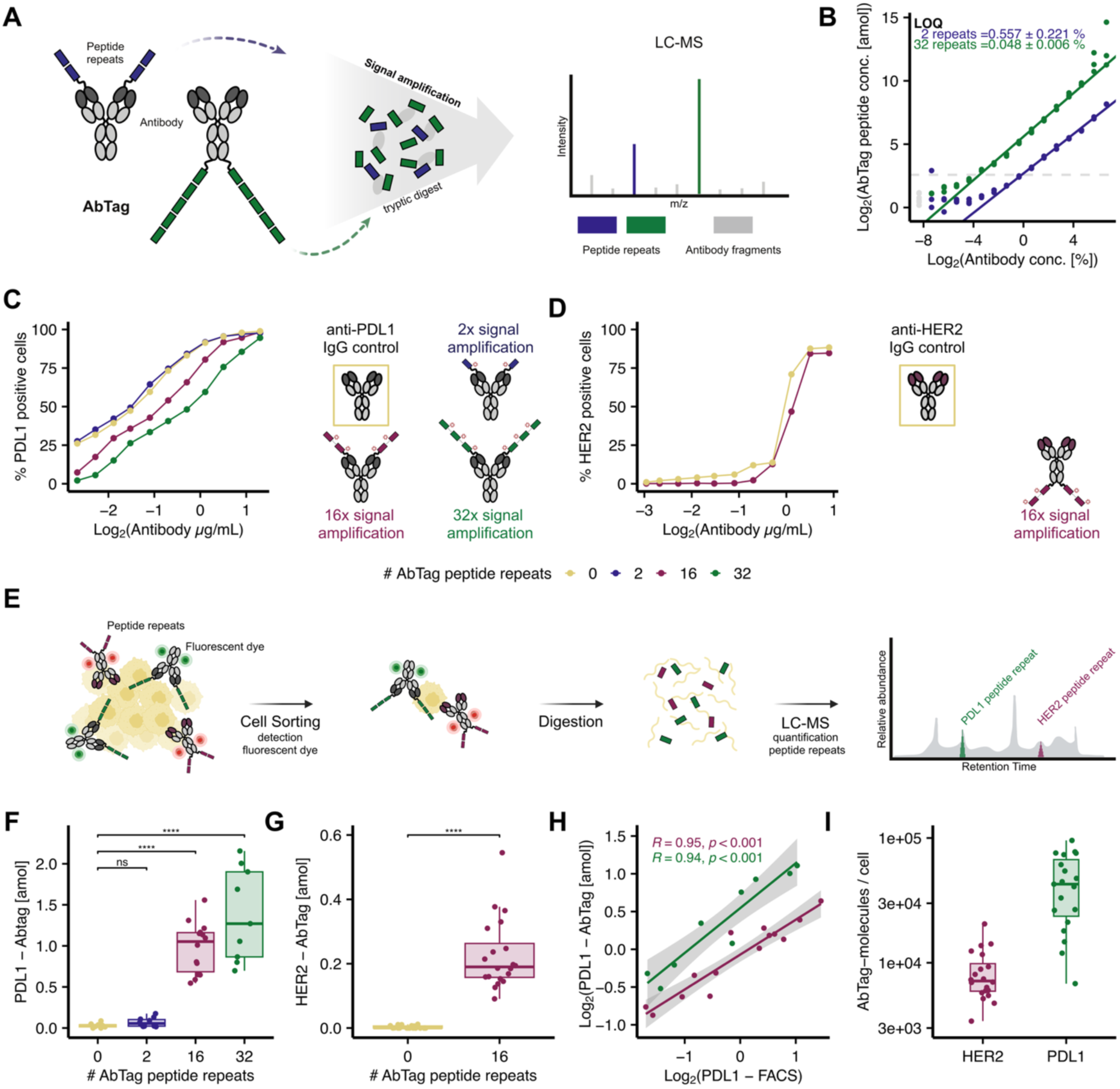
AbTag provides signal amplification through LC-MS detectable tandem-repeat peptides. (**A**) Concept of AbTags highlighting signal amplification. Peptide repeats are conjugated N-or C-terminally to the antibody (**B**) Dilution curve of anti-PDL1 AbTag with 2 and 32 peptide repeats. AbTag peptide repeats quantified by wwPRM using absolute quantification based on stable-isotope-labeled peptides, antibody concentration normalized by antibody-peptides acquired by DIA. Grey line highlights mean + 2 SD around blank. (**C-D**) Mean of triplicates displayed. (**C**) *In vitro* binding characterization for anti-PDL1 AbTags (N-terminal peptide conjugation). PDL1^+^ cell were stained with increasing concentrations of AbTags. anti-PDL1 IgG is shown as positive control. (**D**) *In vitro* binding characterization anti-HER2 AbTags (C-terminal peptide conjugation). HER2^+^ cell were stained with increasing concentrations of AbTags. anti-HER2 IgG as positive control. (**E**) Combined AbTag - scp-MS workflow. (**F-I**) DU-145 cells stained with anti-PDL1 and anti-HER2 AbTag and subjected to scp-MS. AbTag signal quantity of wwPRM-LC-MS and FACS in each single cell displayed. For wwPRM-LC-MS absolute quantification based on stable-isotope-labeled peptides was performed. (**F**) wwPRM-LC-MS quantities of three peptide repeat lengths with anti-PDL1 IgG as negative control tested with pairwise T-test. (**G**) wwPRM-LC-MS quantities of peptide repeats with anti-HER2 IgG as negative control. Significance assessed by Wilcoxon rank sum test. (**H**) Correlation of cell-associated anti-PDL1 AbTag signal quantified by wwPRM-LC-MS and FACS for AbTag constructs with 16 (red) and 32 (green) peptide repeats. (**I**) Number of AbTag molecules bound to single cells based on absolute quantification of AbTag peptide repeats by wwPRM-LC-MS. anti-PDL1 AbTag with 32 peptide repeats (green) and anti-HER2 AbTag with 16 peptide repeats (red) displayed. ns: p > 0.05, * : p < 0.05, ** : p < 0.01, ***: p < 0.001, **** : p < 0.0001

We next asked whether AbTags can sensitively report regulated expression of low-abundance surface proteins while preserving unbiased proteome coverage in a cellular model system. As a proof of principle, we profiled cytokine-induced upregulation of PDL1 in DU-145 cells, combining our anti-PDL1 AbTag with deep scp-MS acquisition (**Fig. 2**). DU-145 cells were treated with IFN-γ, TNF-α or left untreated, stained with an anti-PDL1 AbTag carrying 32 peptide repeats, single-cell sorted and analyzed by DIA scp-MS (**Fig. 2a**). We quantified median 3,275 proteins per cell across three conditions (**Fig. 2b, Supplementary Fig. 3A-C**). While endogenous PDL1 remained undetected, anti-PDL1 AbTag signal showed significant cytokine dependence, with robust but heterogeneous induction in IFN-γ–treated cells and a modest increase after TNF-α–treatment (**Fig. 2c, Supplementary Fig. 3D**). AbTag quantification by LC-MS closely matched the orthogonal readout of the same AbTag measured by flow cytometry across three conditions, confirming that AbTag peptide intensity provides a quantitative proxy for cell-surface PDL1 abundance independent of treatment condition (**Fig. 2D**). This allowed an integration of the AbTag quantified PDL1 response into the global scp-MS proteome (**Supplementary Fig. 3E**).

**Figure 2.**
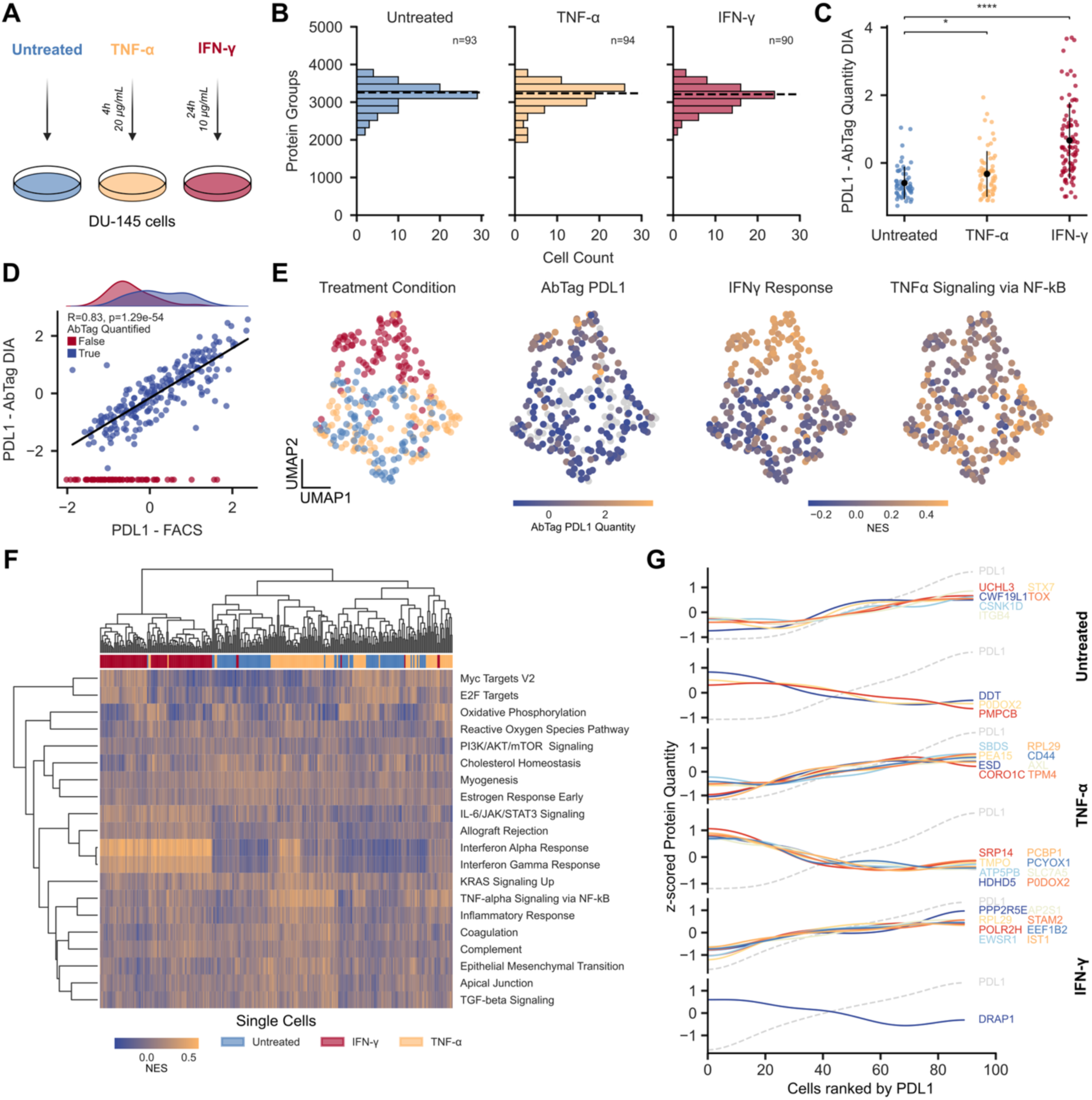
AbTag allows for integration with single-cell proteomics data. **(A)** DU-145 cells were treated with IFN-γ or TNF-α followed by staining with anti-PDL1 AbTag 32-peptide repeats, single-cell sorting and DIA scp-MS acquisition **(B)** Number of proteins obtained per single cell across three conditions. Mean shown as dashed line. **(C)** PDL1 quantity per cell obtained by anti-PDL1 AbTag through DIA-LC-MS. z-scored log-transformed MS1 peptide quantity normalized to cell-size. Mean and standard deviation displayed, pairwise T-test shown. Missing values excluded. **(D)** Correlation of AbTag quantification by DIA-LC-MS (z-scored log-transformed MS1 peptide quantity) and AbTag fluorescence dye (z-scored log-transformed) readout by FACS. Missing values in LC-MS displayed in red. **(E)** UMAP embedding of DU-145 cells based on scp-MS proteome. Treatment, anti-PDL1 AbTag quantity, normalized single-cell pathway enrichment score (NES) for IFN-γ Response and TNF-α Signaling via NF-κB shown. Missing values shown in grey. **(F)** Single-cell pathway enrichment based on MSigDB Hallmarks. 20 pathways with the highest variance between treatments displayed. **(G)** Spearman’s rank correlation for DIA scp-MS proteome with AbTag PDL1 quantity. Top 8 positive/negative correlations with adj p-value below 0.05 displayed as Gaussian kernel. ns: p > 0.05, * : p < 0.05, ** : p < 0.01, ***: p < 0.001, **** : p < 0.0001

To relate AbTag-based PDL1 measurements to global proteome states, we embedded cells based on their endogenous proteomes. UMAP analysis separated IFN-γ treated cells, whereas a modest response to TNF-α was evident due to co-clustering of those cells with control. Importantly, PDL1 AbTag levels co-localized with single-cell enrichment scores for IFN-γ response and TNF-α/NF-κB signaling pathways, thereby confirming IFN-γ and TNF-α induced PDL1 upregulation [8] (**Fig. 2E-F, Supplementary Fig. 4**). Additionally, the endogenous proteome unveiled broad cellular heterogeneity within conditions related to e.g. proliferation. Correlating PDL1 AbTag abundance with the single-cell proteome highlighted both previously described, as well as novel proteins whose levels increase or decrease with PDL1 within each treatment (**Fig. 2G**), including membrane receptors (e.g., CD44, AXL, ITGB4), kinases and phosphatase subunits (CSNK1D, PPP2R5E) and proteins involved in transcription and translation (POLR2H, DRAP1, RPL29) [9], [10]. These data illustrate a proof-of-concept of how AbTags convert an MS undetectable surface marker into a high-dynamic-range, MS-detectable marker that can be mechanistically linked to intracellular proteome remodeling at single-cell resolution.

Together, our results establish a proof-of-principle for AbTag as a general strategy to couple multiplexed, antibody-based detection with unbiased LC-MS proteomics in the same experiment. By encoding antibody targets as tandem-repeat peptide barcodes, AbTags provide signal amplification for low-abundance proteins and allows for absolute quantification. AbTag advances the single-cell proteome below the conventional LOD for endogenous proteins. This sensitivity gain can be achieved with standard LC-MS proteomics workflows, enabling simultaneous targeted and discovery proteomics. The concept can be expanded by combining use case-specific affinity reagents with an effectively unlimited peptide barcode space to achieve target-specific signal amplification in a highly multiplexed fashion.

## Methods

### Selection of antibodies and non-human peptides

Two non-human peptides (TASEFDSAIAQDK and LTILEELR) were selected for this study after being run through PEP-FOLD4 to assess their folding configuration. The correspondent DNA sequences were ordered from GeneWiz. Anti-PDL1 and anti-HER2 antibodies were selected for targeting PDL1 and HER2 epitopes, respectively, and served as backbones for the attachment of non-human peptides. The anti-PDL1 antibody incorporated the variable (VH-VL) domain of atezolizumab, while the anti-HER2 antibody the VH-VL domains of trastuzumab (Herceptin). In both constructs, the constant Fc region was derived from rituximab. Peptide repeats were added N-terminally to anti-PDL1 antibodies and C-terminally to anti-HER2 antibodies using a flexible 10-mer GS linker (GGGGS)_2_.

### Primer design

The primers used for plasmid linearization and insert amplification were designed using the online software Benchling. All primers were formed by 15 to 40bp, 40% to 60% GC content and had a melting temperature (Tm) ranging from 65°C to 75°C. When required, an overlap exceeding 12bp between parental and template strains was included.

### Plasmid linearization and inserts amplification through PCR reaction

All vectors used in the study originated from the pcDNA3.1 vector backbone containing antibody constant domains (IgG1 for heavy chain and Cκ for light chain) either targeting PDL1 or HER2 [6]. Plasmid linearization and insert amplification were performed using PCR reaction.

Plasmids were linearized in a PCR mixture, using Q5® Hot Start High-Fidelity 2X Master Mix (NEB, #M0494S), forward- and reverse primers, and 10 ng of template DNA. PCR was performed such as an initial denaturation step at 98 °C 30 s, followed by 30 cycles utilizing the following parameters: denaturation at 98 °C for 10 s, annealing at 72 °C for 10 s, and extension at 72 °C or 68°C for 3 mins and 10 s, respectively. Subsequently, a final extension step at 72 °C for 5 min was performed. Similarly, inserts containing the genes of interest were amplified in a PCR mixture, as described before. Nevertheless, extension was performed at 72 °C for 30 s and a final extension step at 72 °C for 2 min.

DNA isolation of the PCR product was achieved using the GeneJet Gel Extraction Kit (Thermo Scientific, #K0691), followed by Monarch^®^ PCR & DNA Cleanup Kit (NEB, #T1030). The integrity of the isolated DNA was assessed through 1% agarose gel electrophoresis and the NanoDrop spectrophotometer (ThermoFisher Scientific).

### HIFI cloning

Linearized plasmids and amplified inserts were ligated using NEBuilder^®^ HiFi DNA Assembly Cloning Kit (NEB, #E5520S). A 2:1 insert:vector ratio was used with a final concentration of 0.15 pmol of DNA. The HIFI reaction totaled 20 μL, with an insert and plasmid volume varying for each mixture, 10 μL of NEBuilder^®^ HiFi DNA Assembly Master Mix and miliQ water to the set final volume. The mixtures were incubated at 50 °C during 15 mins, in a Thermocycler (Eppendorf® Mastercycler®).

### Transformation into DH5α *E. coli* cells

HIFI products underwent transformation into NEB^®^ DH5α Competent *E. coli* (NEB, #C2987H) right after incubation, following manufacturer instructions. The recombinant plasmids were confirmed by sequencing (Macrogene, Amsterdam, Netherlands) and whole plasmid sequencing (UnveilBio, Copenhagen, Denmark). Plasmids confirmed with desired insertions were purified using NucleoBond Xtra Midi Plus EF (Macherey-Nagel) to ensure endotoxin removal.

### Transient production of antibodies in HEK293 cells

All antibodies were produced by transient expression in HEK293F cells, aiming for secretion into the culture supernatant. Plasmids encoding either heavy chain (HC) or light chain (LC) were transiently cotransfected using linear polyethylenimine (PEIPro, Polyplus-transfection) according to manufacturer instructions and transfected cells were cultured for 7 days in FreeStyle 293 Expression Medium (12338026, Gibco) at 37 °C and 5% CO_2_. A 3:2 transfection ratio (HC:LC) was employed for all antibodies. After incubation, the supernatants were clarified (centrifugation at 3000 ×g for 10 min followed by 1500×g for 10 min and 0.22 μm filtering) before purification using Protein A (HiTrap MabSelect SuRe, Cytiva, lot: 10293990) affinity chromatography connected to an ÄKTA Pure system (Cytiva). All antibodies were concentrated, and buffer was exchanged to phosphate-buffered saline (PBS) using Amicon Ultra-15 centrifugal filter devices (30-kDa cutoff, EMD). The protein concentrations were measured using the NanoDrop One UV-Vis spectrophotometer (ThermoFisher). The quality of the purified antibodies was analyzed by SDSPAGE (4–20% Mini-PROTEAN® TGX™ Precast Protein Gels, Bio-Rad) in the presence and absence of 25 mM 1,4-dithiothreitol (DTT) and visualized through Coomassie Brilliant Blue staining.

### Basic cell line culture and maintenance

Human cell lines Raji PDL1^+^ [11], HT29 and DU-145 were maintained in complete RPMI (RPMI 1640 (Sigma-Aldrich) supplemented with 10% FBS (Gibco), 1% L-glutamine and 1% penicillin-streptomycin (Sigma-Aldrich)). Adherent cell lines (HT29 and DU-145) were split twice weekly when approaching 90% confluency. Raji PDL1^+^ were passaged every 2-3 days and seeded at 0.5*10^6^ cells/mL. All cell lines were kept at 37 °C in a humidified incubator with 5% CO_2_. In experiments that involved cytokine treatment DU-145 cells were stimulated with either IFN-γ (24h 10 µg/mL) or TNF-α (4h 20 µg/mL) at 37 °C.

### Antibody binding assessment via flow cytometry

Antibodies were labelled according to manufacturer protocol with Zenon™ Human IgG Labeling Kit (Thermo Fisher Scientific, #Z25402) (Thermo Fisher Scientific, #Z25408). Cells were stained with LIVE/DEAD^TM^ Fixable Yellow Dead Cell Stain (ThermoFisher Scientific, #L34959) for detection of dead cells and incubated for 10min with FcR blocking reagent (Miltenyi, #130059901). Raji PDL1^+^ and HT29 were incubated for 10 min on ice with labelled antibodies (0-20μg/mL), washed and fixed with fixation buffer (eBioscienceTM, #88-8824-00) prior to analysis on the MACSQuant Analyzer 16 flow cytometer (Miltenyi). Data from three independent assays using cells from different passages was analyzed in Prism 10.2.1 (GraphPad Software) where data was fitted and EC50 values were calculated.

### Fluorescence activated cell sorting (FACS) for scp-MS analysis

Prior to FACS, DU-145 cells were washed in DPBS, and antibodies and AbTags were labelled according to manufacturer protocol with Zenon™ Human IgG Labeling Kit (Thermo Fisher Scientific, #Z25402 or Thermo Fisher Scientific, #Z25408). Following this, 1*10^6^ cells were incubated with labelled antibodies or AbTags for 10 min on ice. Cells were washed in DPBS to eliminate excess antibody before sorting on a Sony Sorter MA900 cell sorter using a 130 μm sorting chip. Doublets and cell debris were excluded based on forward-and side scatter. For experiments shown in Figure 1 PDL1 and HER2 positive populations were gated using unstained controls. Cells were sorted in a 384 well plate containing lysis buffer as described below.

### Proteomics sample preparation

Single cell proteomic sample preparation was performed in a 384 well plate as described previously [7]. Briefly, 1uL 20 % 2,2,2-Trifluoroethanol (TFE, Sigma-Aldrich, 91683) with 80 mM Triethylammonium Bicarbonate (TEAB, 90114 Thermo Scientific) was added to each well of a 384 Eppendorf™ twin.tec™ PCR Plates LoBind™. Cells were sorted as described above and the plate was snap frozen on dry ice after sort was completed. Cells were heated up to 96 °C for 5 min followed by freezing to -80 °C. 1uL of 100 mM TEAB containing 2 ng Trypsin Platinum (VA9000, Promega, USA) was added to each well and the plate was incubated overnight (ON) at 37 °C. The digestion was quenched with 1 uL 1 % TFA. If the experiment included isotope labeled peptides, these were added within the 1 % TFA. Stable isotope labeled (SIL) peptides were added at a concentration of 1.56 amol/cell (Pierce® Peptide Retention Time Calibration Mixture PN 88320, 0.5pmol/µL). The same protocol was used to assess digestion reproducibility of the anti-PDL1 AbTags. Here 500 amol SIL peptides were added to each digest.

Assessment of the quantitative performance of the AbTags was performed in a stable 100 pg/uL Pierce™ HeLa Protein Digest Standard. SIL peptides were added at a concentration of 1.56 amol/uL as stated above. The AbTag was previously heated to 95 °C for 10 min followed by reduction in 2% sodium deoxycholate (SDC), 1x Phosphate-buffered saline (PBS), 10 mM Tris(2-carboxyethyl)phosphine (TCEP) for 30min at 37 °C. The sample was alkylated at room temperature for 30 min in 40 mM chloroacetamide (CAA) before it was diluted to 0.1 % SDC. Trypsin was added in a 1:25 ratio and the sample was digested at 37 °C ON. After quenching to 0.5 % TFA the sample was desalted using the SOLAμ™ SPE Plates (Thermo Scientific™). The AbTag digest was diluted in the previously prepared 100 pg/uL hela digest.

### LC-MS analysis

Experiments displayed in Figure 1 were performed on an Ultimate 3000 RSLCnano system with µPAC NEO low load Trapping Column (Thermo Scientific) and µPAC NEO HPLC Column 50 cm (Thermo Scientific). The column was kept at 50 °C and the autosampler was operated at 7 °C. Samples were loaded at 750 nL/min in 99 % Solvent A (1% Formic Acid). Solvent B (80% Acetonitrile, 1% Formic Acid) was increased to 8 % for 0.2 min followed by a drop in flow to 200 nL/min and a linear gradient to 24% Solvent B over 2.8min. Solvent B was increased to 48 % until minute 8 followed by an increase to 99 % B for 6.55 min. The column was equilibrated for the remaining 2.85 min and the next precolumn loading at 1 % B with a flowrate of 750 nL/min. Experiments shown in figure 2 were performed on a Vanquish™ Neo (Thermo Scientific) with a PepMap™ Neo Trap Cartridge precolumn and an Ion Optics Aurora® Rapid™ 8×75 XT C18 UHPLC column. The column was operated at 50 °C and the autosampler at 7 °C. Samples were loaded at 800 nL/min in 96 % Solvent A. Solvent B was increased to 10 % for 0.75 min, 16 % at min 0.263, 33 % at 1.063 min and to 99 % B at 1.113 min. The flow was dropped to 100 nL/min until minute 7.1 min. The flow was increased again to 800 nL/min and at minute 7.1 Solvent B was reduced to 4 %.

Experiments related to Figure 1 were performed on an Orbitrap Eclipse mass spectrometer connected to FAIMS Pro operating at -45 V for DIA experiments. MS1 acquisitions were performed at 120k resolution with an AGC target of 300 % and an injection time (IT) 246ms.

MS2 scans operated at a HCD of 26% with a resolution of 120k with 246 ms IT. The AGC target was set to 1000 % and 6 isolation windows covering 400 – 800 m/z were used. Targeted MS2 scans were performed at 240k resolution with 504 ms IT, an AGC target of 300 %. Both FAIMS CVs and NCE parameters were optimized for each target. Precursors were isolated at a 1.7, 5.7 or 6.7 m/z window to isolate both endogenous and isotopically labeled peptides as previously described [7]. PRM and DIA scans were combined in one method where PRM scans operated in a 0.7 min retention time window.

Experiments in Figure 2 were performed using an Orbitrap Astral mass spectrometer with FAIMS Duo Pro. The FAIMS unit was operated at -48 V and samples were analyzed in DIA. MS1 scans were performed at 240k resolution with an injection time of 100 ms and an AGC target of 500 %. MS2 scans were performed at a HCD of 25 % with an AGC target of 500 % over the mass range of 150-2000 m/z. Ions were injected for 40 ms over 20 variable MS2 windows spanning from 400-800. The loop control was set to N = 13.

## Data Analysis

PRM and wwPRM results were analyzed in Skyline-daily (version 25.1.1.206). A target library was built on the SIL peptides. All y-ions were used for peptide identification and peak area integration. A report at fragment level peak area was exported and further analysis was performed in R. The top 3 y-ions (TASEFDSAIAQDK - y9,y11,y8 and LTILEELR - y6,y5,y4) of each heavy and light peptide were selected and summed for quantification. The light peptide peak area was divided by the heavy peptide peak area for normalization and determination of the molar light peptide peak concentration. To determine the amount of antibody molecules that are bound to each cell we determine the number of light peptide molecules per well and divide it by the number of peptide tags coupled to each antibody:

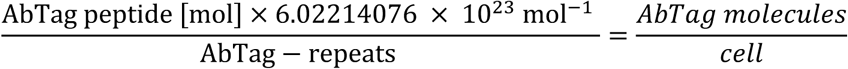

In addition to the wwPRM data, DIA spectra were searched with DIA-NN 2.1.0 for all single cell experiments. Samples were searched without MBR in a first search to exclude dropouts and searched with MBR after filtering. Maximum number of missed cleavages was set to 1 with trypsin/P as digestion enzyme. Both N-terminal M-excision and cysteine carbamidomethylation were included. Protein interferences were set to genes and precursor level FDR was set to 1 %. AbTag dilution curves were searched with Spectronaut 19.0. Settings were kept at default with the changes of quantification level set to MS1, fixed modifications were removed. Data was exported at precursor level. In all cases the Uniprot reviewed human proteome (2022) was used to build the predicted library. AbTag peptides and antibody backbone sequences were added to the FASTA.

The wwPRM and DIA data was combined with index data from the FACS sort. For figure 1 cells were included based on >1000 protein groups and a robust linear regression fitted between the FSC.A and the number of precursors for all cells. The PDL1 and HER2 fluorophore dye abundance was selected to be below the maximum 0-peptide repeat AbTag signal to ensure even cell distribution. For figure 2 cells were filtered based on the following criteria: the average full width half max lies within mean ± standard deviation (sd) of the dataset, the number of identified precursors and total MS1 signal lies within mean ± 2 sd of the dataset. Cells follow a linear relationship of FSC.A vs nr. precursor. Furthermore, cells with a protein level data completeness of <50% are excluded. Leiden clustering was used to identify technical outliers. Single-cell data was imputed to have half the minimal quantity of a quantified protein if the protein was identified in at least 30 % of all cells. Protein expression for each protein was median normalized and log transformed.

Standard curves of 2-and 32-peptide repeat AbTags were calculated based on all dilution points with a CV of <20 % between replicates. To ensure equal concentrations of antibody backbone between conditions, the peptide repeat signal was normalized to the peptides identified from the antibody backbone. The average of 2, 4 and 6 sd around the blank signal was defined as the smallest theoretically possible limit of quantification (LOQ). The ratio between all remaining data points was taken and compared to the expected ratio. Ratios between dilution points spanning up to an 8-fold distance were compared. Dilution-Ratio-Pairs with a deviation of the expected ratio larger than 25 % were excluded and a linear weighted model was fitted to the remaining data points. Based on this regression the LOQ was calculated as the last point at which the linear fit can be distinguished from a lower dilution point. All data analysis was performed in R and python.

## Data availability

All MS raw files, database search results, and Skyline files are available through Panorama Public with the ProteomeXchange identifier PXD072099.

## Code availability

Data to generate the figures in the manuscript are available through Panorama Public. Code used for analysis is available on Github and through Panorama Public.

## Competing interests

A patent application for MS-detectable peptide tags has been filed by the Technical University of Denmark (DTU). The Schoof lab has a sponsored research agreement with Thermo Fisher Scientific, the manufacturer of the instrumentation used in this research. However, analytical techniques were selected and performed independently of Thermo Fisher Scientific. The remaining authors declare no competing interests.

## Acknowledgments

We thank Valdemaras Petrosius for valuable feedback on the manuscript. The DU-145 cell line was kindly gifted by Per Thor Straten from Centre for Cancer Immune Therapy (CCIT). This work was funded by the following grants to E.M.S.: 1) reference number NNF21OC0071016 from the Novo Nordisk Foundation; 2) case no. 2067-00053B from the Independent Research Fund Denmark, 3) Lundbeck Foundation (R413-2022-869), and 4) Velux Foundation (00053026). Work in the Porse lab was supported by grants from the Svend Andersen Foundation, the Eva and Henry Frænkel Memorial Foundation and the Independent Research Fund Denmark. S.G. gratefully acknowledges the Novo Nordisk Foundation for funding this work through the following grants: NNF20SA0066621, NNF19SA0056783, NNF21SA0072683, and NNF19SA0035474.

---

**Supplementary Figure 1.**
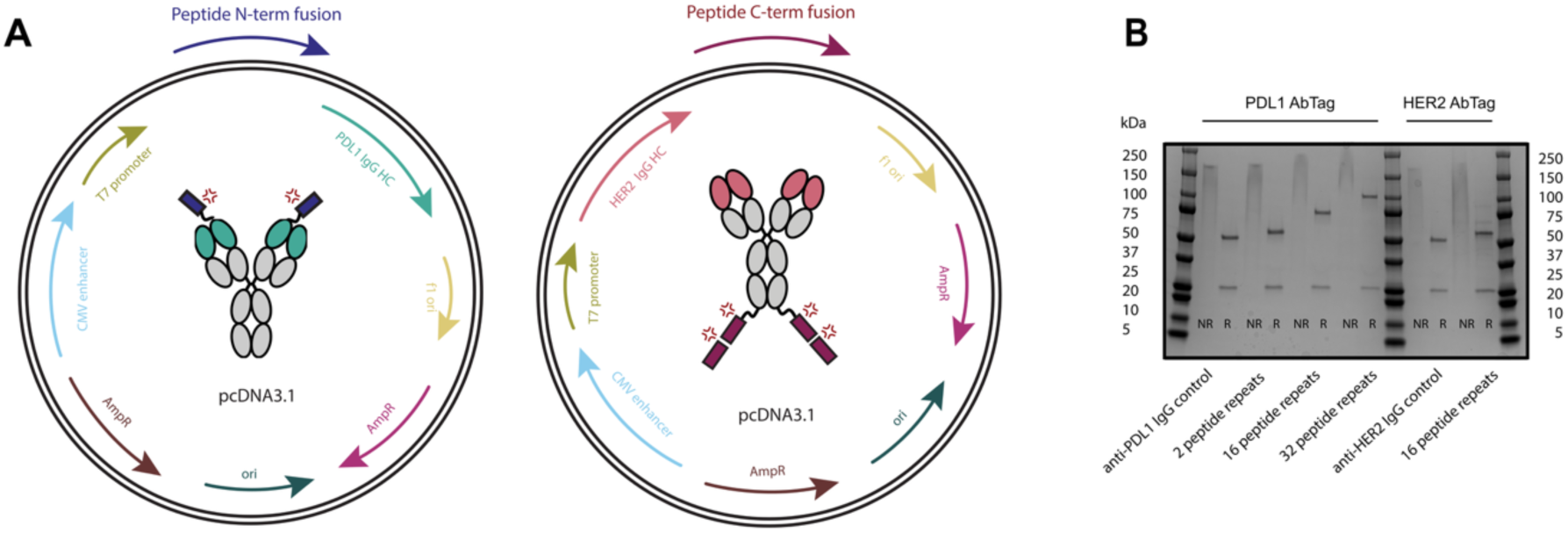
Overview of the AbTag design. **(A)** pcDNA3.1 vectors including the insertion site for the peptide of interest. **(B)** SDS-PAGE gel confirming HC and LC sizes. Ladder is in left and right lanes, reduced (R) and non-reduced (NR) samples indicate appropriate size of the AbTags. Anti-PDL1 and anti-HER2 IgG antibodies were used as controls, underlining HC size increase due to the peptide tags.

**Supplementary Figure 2.**
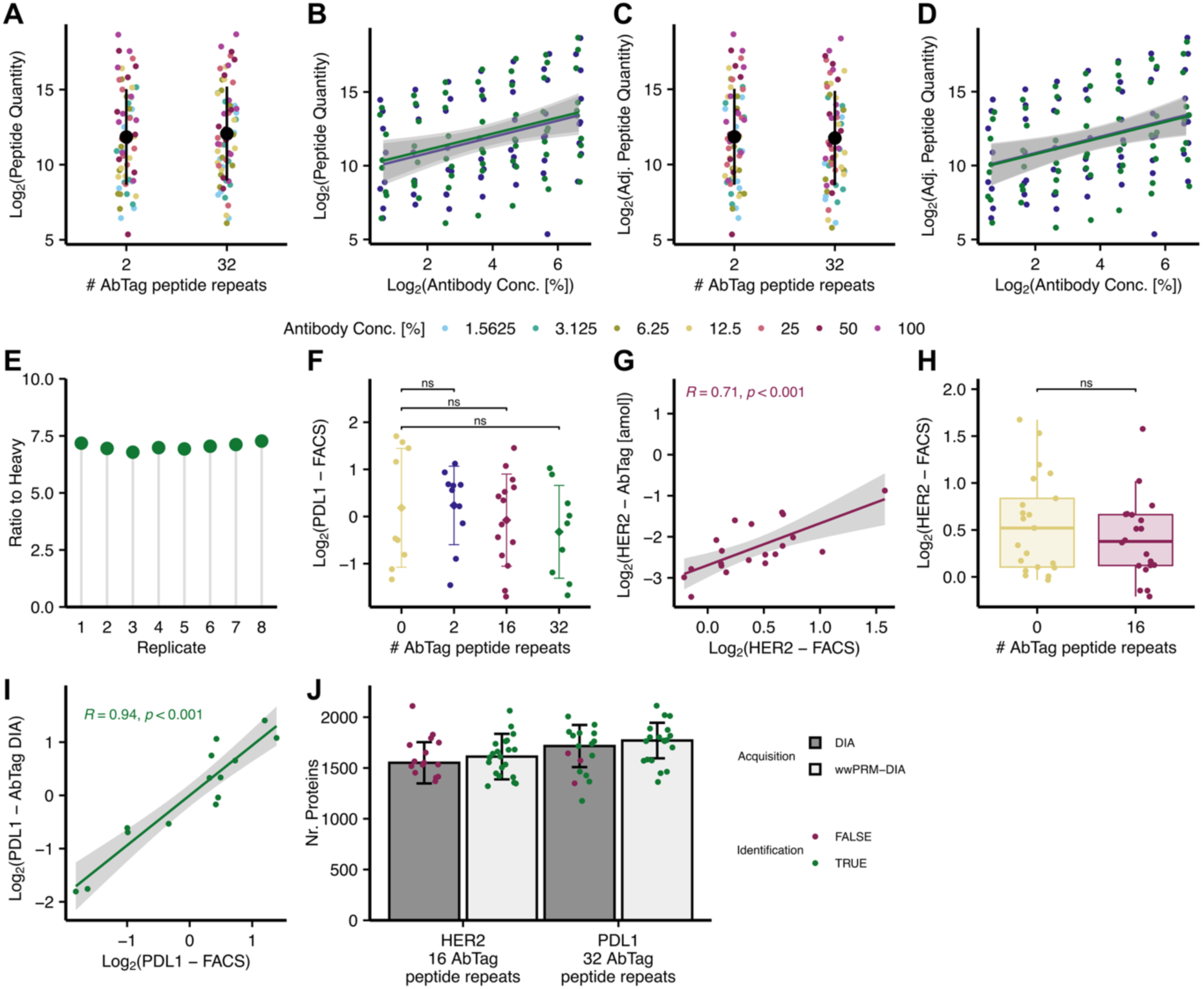
AbTag LC-MS detection and quantitative performance (A-D) Normalization of antibody concentration between AbTags with 2- and 32 peptide repeats shown in figure 1B. (**A**) Antibody peptides quantified before normalization. (**B**) Antibody peptides quantified in the top 7 dilution points of the dilution curve. (**C**) Antibody peptides quantification normalized between AbTags. **(D)** Antibody peptides quantified in the top 7 dilution points of the dilution curve after normalization between AbTags. **(E)** Repeated digestion of AbTag with 32-peptide repeats. Quantification based on PRM acquisition in ratio of AbTag peptide repeats to SIS peptide (Ratio to Heavy, CV = 2.2 %) (**F**) PDL1 abundance on cells based on Alexa 488 fluorescence detected during FACS sorting. (**G**) Correlation of HER2 abundance detected on cells with Alexa 647 fluorescence determined by FACS and AbTag quantification by wwPRM-LC-MS. **(H)** HER2 abundance on cells based on Alexa 647 fluorescence detected during FACS sorting. **(I)** Correlation of PDL1 abundance detected on cells with Alexa 488 fluorescence determined by FACS and AbTag quantification by DIA-LC-MS. **(J)** Number of proteins identified per cell during LC-MS acquisition with DIA or wwPRM-DIA searched with DIANN 2.1.0. Color displays whether anti-HER2-16-AbTag peptide and anti-PDL1-32-AbTag peptide could be quantified with the respective method. Pairwise-T-test corrected by fdr. ns: p > 0.05, * : p < 0.05, ** : p < 0.01, ***: p < 0.001, **** : p < 0.0001

**Supplementary Figure 3.**
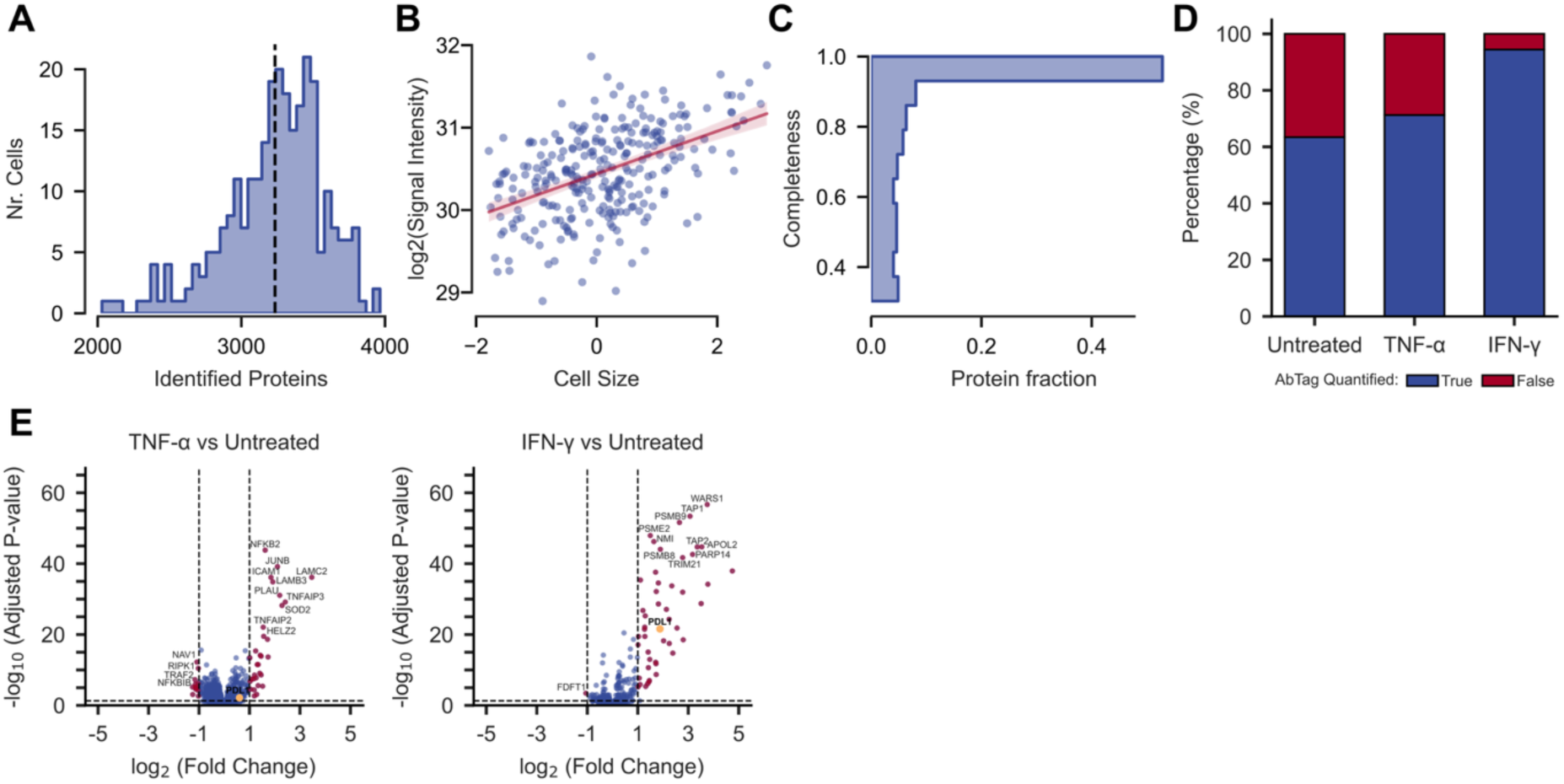
AbTag integration with single-cell proteomics data. **(A)** Overall identified number of proteins by scp-MS across treatment conditions shown in figure 2 **(B)** log2 MS1-signal of each single cell correlated with the cell size (z-scored FSC.A) determined by FACS. **(C)** Data completeness across identified proteins. **(D)** Percentage of single cells in which PDL1 AbTag peptide was quantified by DIA LC-MS grouped by treatment. **(E)** Differential expression analysis of IFN-γ and TNF-α vs. untreated control. PDL1 quantified by AbTag shown in yellow.

**Supplementary Figure 4.**
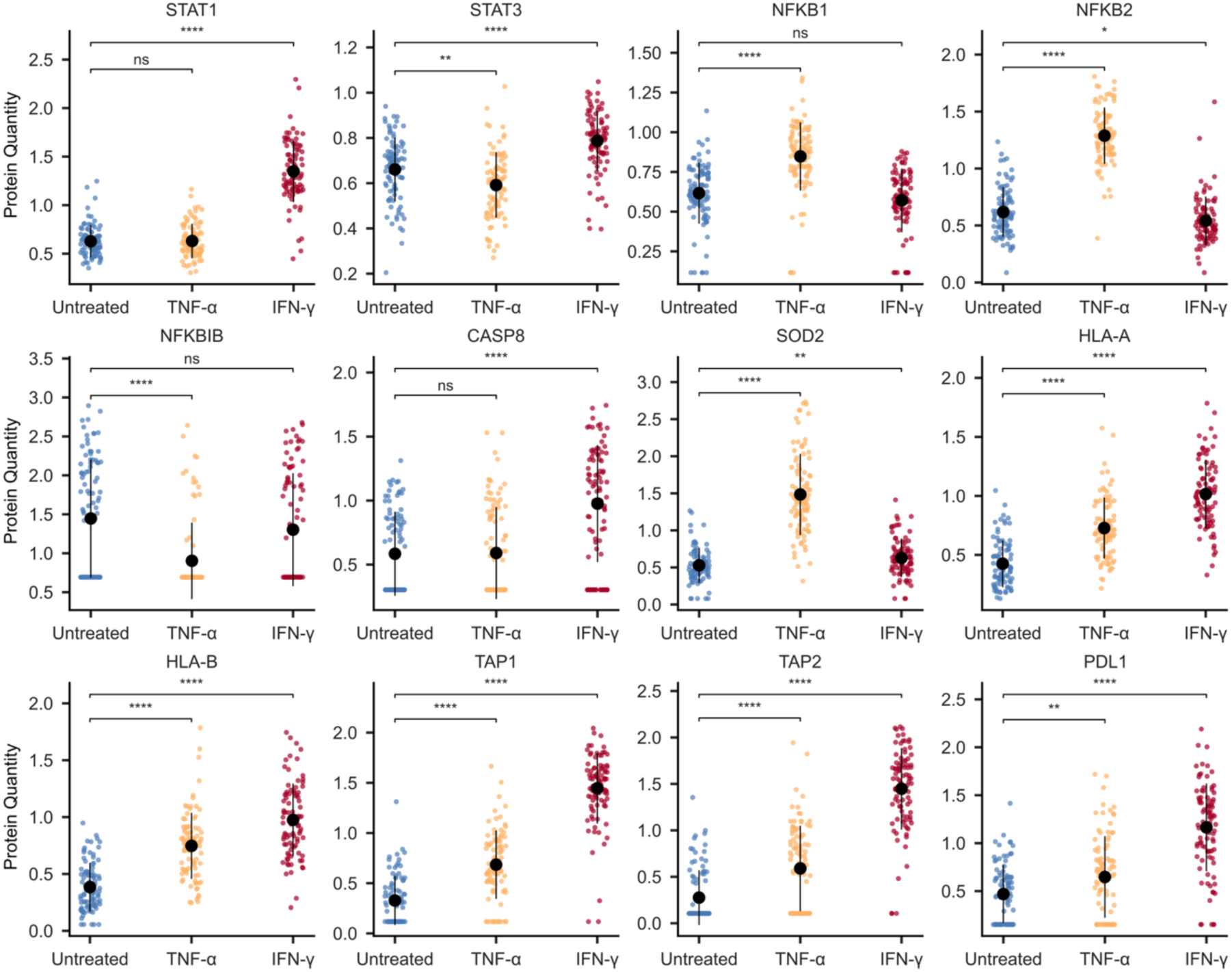
: scp-MS captures known cellular responses to cytokine treatment. Log-transformed median normalized protein quantities determined by scp-MS for selected example proteins involved in cellular processes related to treatment conditions. PDL1 quantified by anti-PDL1 AbTag. Imputed protein quantities displayed as half of the minimum measurement. Mean and standard deviation displayed. Pairwise T-test of untreated control to treatment conditions. ns: p > 0.05, * : p < 0.05, ** : p < 0.01, ***: p < 0.001, **** : p < 0.0001

## References

[1] B. Budnik, E. Levy, G. Harmange, and N. Slavov, “SCoPE-MS: mass spectrometry of single mammalian cells quantifies proteome heterogeneity during cell differentiation,” Genome Biol., vol. 19, no. 1, p. 161, Oct. 2018, doi: 10.1186/s13059-018-1547-5.

[2] V. Petrosius et al., “Quantitative Label-Free Single-Cell Proteomics on the Orbitrap Astral MS,” Mol. Cell. Proteomics, p. 100982, May 2025, doi: 10.1016/j.mcpro.2025.100982.

[3] D. R. Bandura et al., “Mass Cytometry: Technique for Real Time Single Cell Multitarget Immunoassay Based on Inductively Coupled Plasma Time-of-Flight Mass Spectrometry,” Anal. Chem., vol. 81, no. 16, pp. 6813–6822, Aug. 2009, doi: 10.1021/ac901049w.

[4] L. M. Park, J. Lannigan, and M. C. Jaimes, “OMIP-069: Forty-Color Full Spectrum Flow Cytometry Panel for Deep Immunophenotyping of Major Cell Subsets in Human Peripheral Blood,” Cytometry A, vol. 97, no. 10, pp. 1044–1051, 2020, doi: 10.1002/cyto.a.24213.

[5] M. Stoeckius et al., “Simultaneous epitope and transcriptome measurement in single cells,” Nat. Methods, vol. 14, no. 9, pp. 865–868, Sept. 2017, doi: 10.1038/nmeth.4380.

[6] A. V. Madsen, P. Kristensen, A. K. Buell, and S. Goletz, “Generation of robust bispecific antibodies through fusion of single-domain antibodies on IgG scaffolds: a comprehensive comparison of formats,” mAbs, vol. 15, no. 1, p. 2189432, Dec. 2023, doi: 10.1080/19420862.2023.2189432.

[7] J. Woessmann, et al., “Informed Data-Independent Acquisition Enables Targeted Quantification of Key Regulatory Proteins in Cell Fate Decision at Single-Cell Resolution,” June 02, 2025, bioRxiv. doi: 10.1101/2025.05.30.656945.

[8] D. Korentzelos, A. Wells, and A. M. Clark, “Interferon-γ increases sensitivity to chemotherapy and provides immunotherapy targets in models of metastatic castration-resistant prostate cancer,” Sci. Rep., vol. 12, no. 1, p. 6657, Apr. 2022, doi: 10.1038/s41598-022-10724-9.

[9] T. Kong et al., “CD44 Promotes PD-L1 Expression and Its Tumor-Intrinsic Function in Breast and Lung Cancers,” Cancer Res., vol. 80, no. 3, pp. 444–457, Feb. 2020, doi: 10.1158/0008-5472.CAN-19-1108.

[10] H. D. Skinner et al., “Integrative Analysis Identifies a Novel AXL–PI3 Kinase–PD-L1 Signaling Axis Associated with Radiation Resistance in Head and Neck Cancer,” Clin. Cancer Res., vol. 23, no. 11, pp. 2713–2722, May 2017, doi: 10.1158/1078-0432.CCR-16-2586.

[11] E. Hui et al., “T cell costimulatory receptor CD28 is a primary target for PD-1-mediated inhibition,” Science, vol. 355, no. 6332, pp. 1428–1433, Mar. 2017, doi: 10.1126/science.aaf1292.

