## Supplementary figures and images for "Antibody-based signal amplification for single-cell proteomics by mass spectrometry"

### Supplementary Figure 1

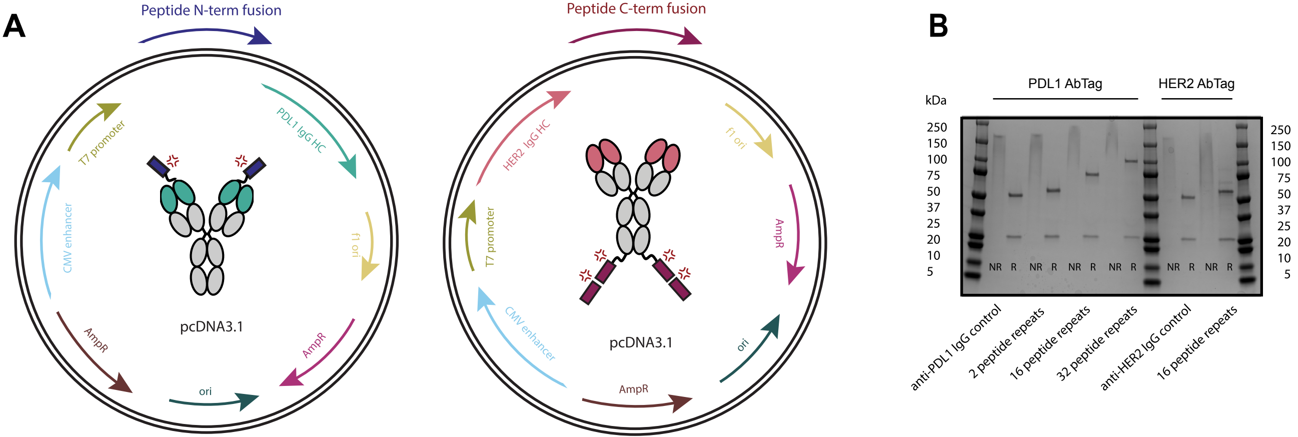

### Supplementary Figure 2

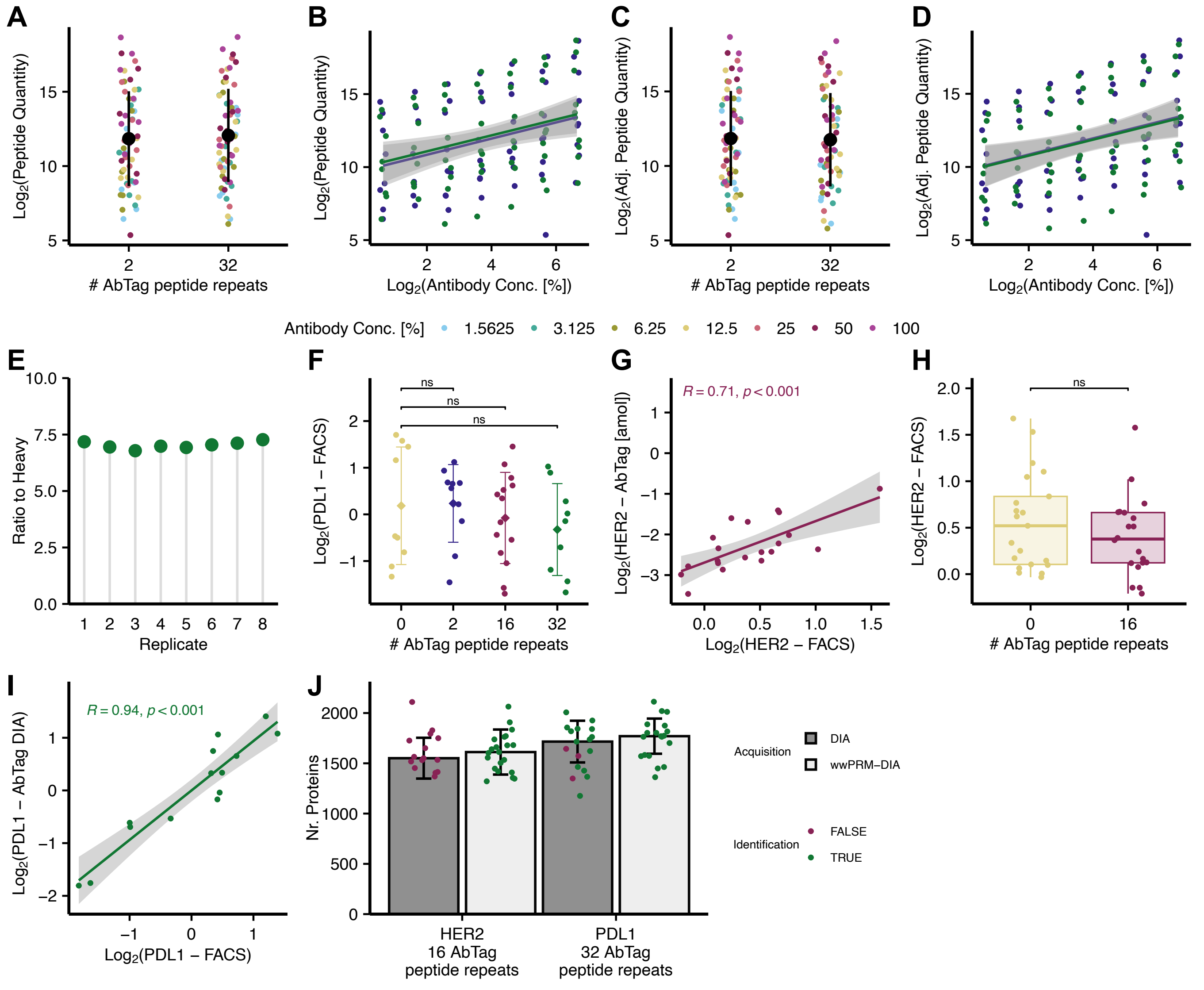

### Supplementary Figure 3

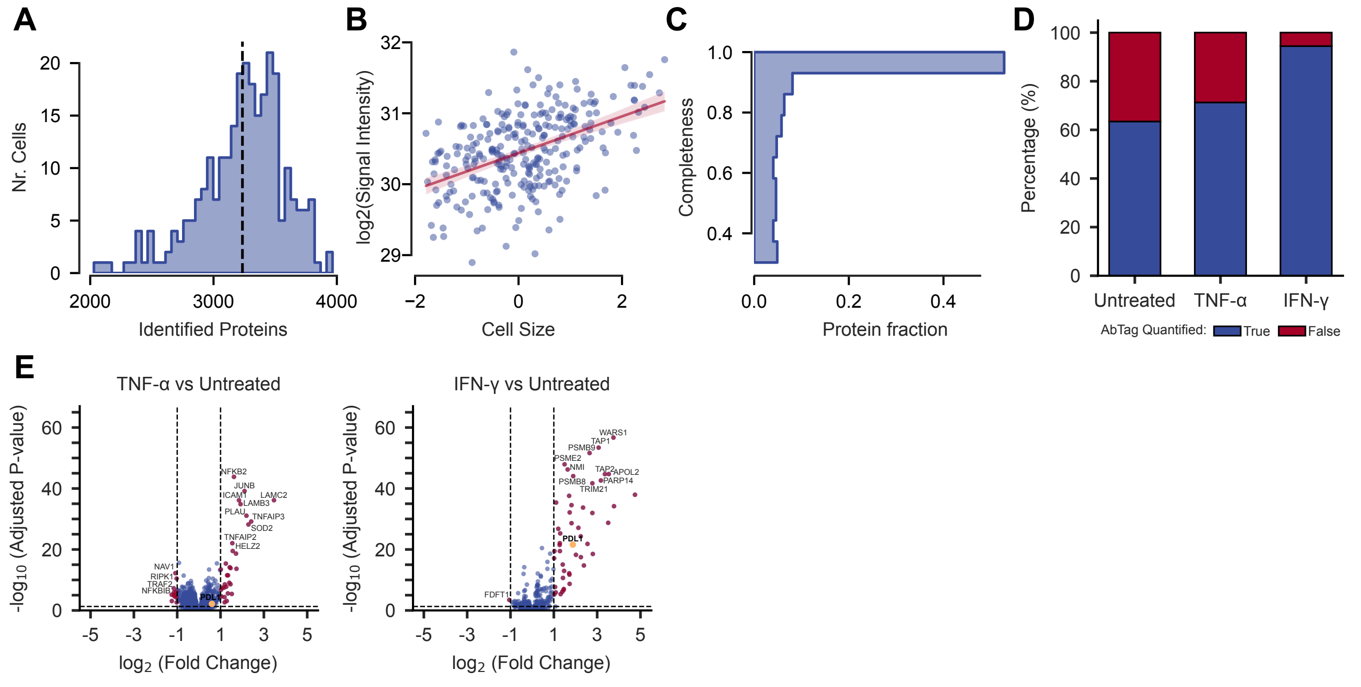

### Supplementary Figure 4

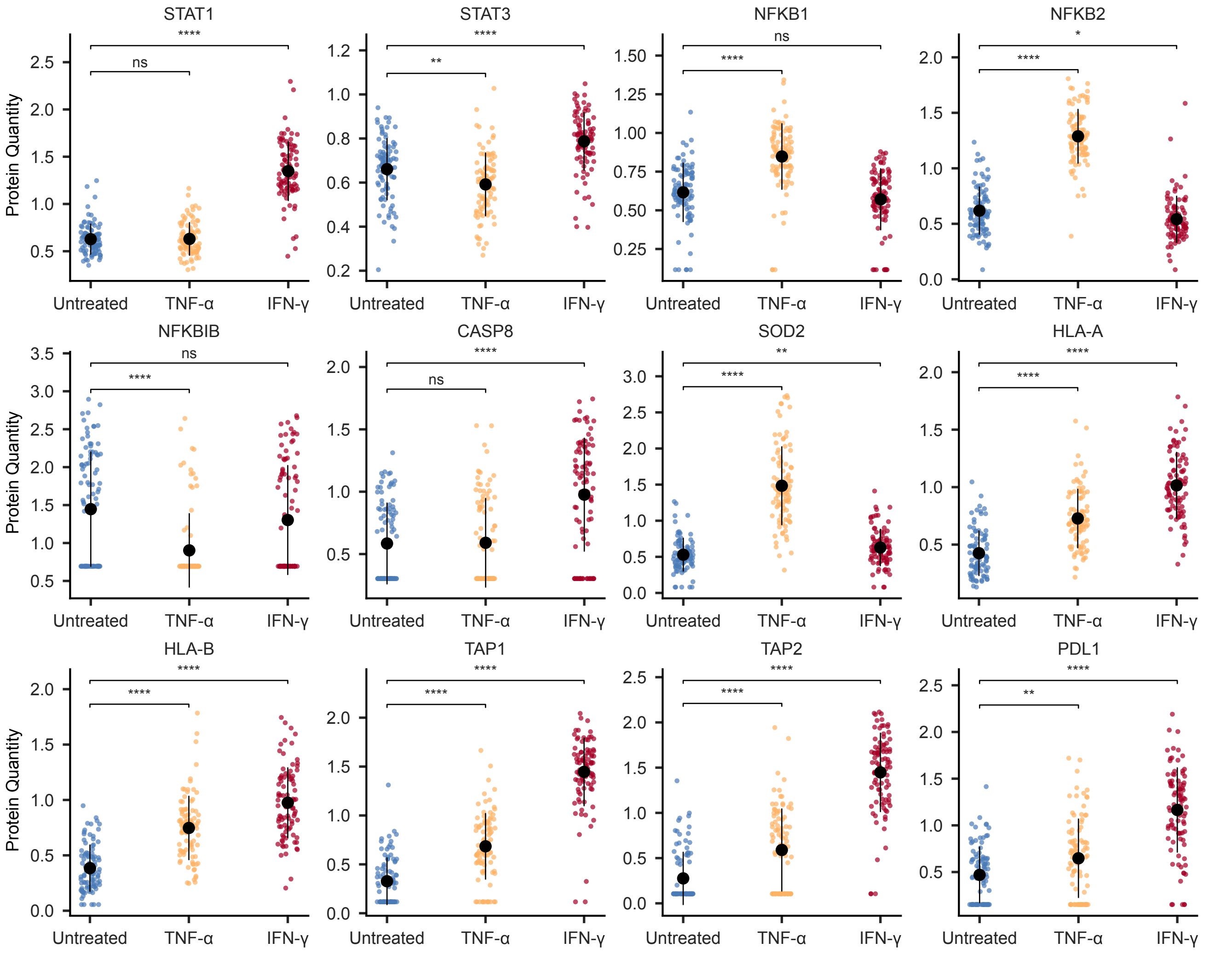
